# Novel mark recapture surveys reveal stable pigmentation patterns but age-related melanism in the common frog *Rana temporaria*

**DOI:** 10.1101/2024.05.11.593487

**Authors:** Silviu O. Petrovan, Sharvani Sivakumar, Carissa Li, Alec Christie

## Abstract

Variations in melanin-based pigmentation patterns are frequent evolutionary adaptations in vertebrates and allow the study of phenotypic diversity across populations as well as individual recognition. Age-related changes in such pigmentation patterns have been demonstrated for several species but remain rarely explored in adult amphibians. For the wide-ranging European common frog (*Rana temporaria*) dorsal melanism was discussed in the context of UV protection at high altitude or as thermal adaptation at high latitude, yet patterns remain unquantified across most of its range or in terms of individual variation over time. We investigated dorsal pigmentation patterns in a wild population of *R. temporaria* in a lowland site in England which was the focus of a 7 year high-intensity mark recapture programme. We collected dorsal photographs using phone cameras and analysed pigmentation patterns using visual classifications and ImageJ image processing software. Pigmentation was highly variable between individuals but overall: (i) the number of melanin-based spots was largely stable or decreased over time as enlarging spots sometimes merged but never disappeared; (ii) the total melanin-based pigmented area increased significantly over time on average, albeit not for all individuals; and (iii) pigmentation was similar in males compared to females although the darkest individuals tended to be males. Our analysis reveals important patterns in pigmentation change and stability in this widespread species but the mechanisms and specific drivers remain unknown.

## Introduction

Melanin-based colour polymorphisms are common evolutionary adaptations across animal groups (birds: Gangoso et al., 2011; fish: Kittilsen et al., 2009; insects: Ethier et al., 2015; reptiles: Ducrest et al., 2014). Consequently, the study of melanin patterning within and between species and populations has been repeatedly employed to understand the links between ecology and evolution. Melanin plays a role in various non-mutually exclusive adaptive functions in animals, linked to different aspects of life history. For example, it is associated with preferred prey specialisation (Karell et al., 2021), protection against UV radiation (Roulin, 2014), thermoregulation (Vences et al., 2002), resistance to parasitism (Jacquin et al., 2011), as well as pleiotropic effects linked to phenotypically plastic traits such as immunity, body mass and hormone levels (Roulin, 2016). Melanin also has a putative role in defence against predators by crypsis mediated by light sensitivity (Fulgione et al., 2014) or by aposematism (Turner, 1977).

Intraspecific variation in melanic pigmentation can manifest as either distinct colour morphs (King, 1987) or along a spectrum of varying degrees of dark pigmentation in individuals (Riobo et al., 1999). Mechanisms for changing pigmentation patterns during the lifespan of the individual can be split into: (i) short-term physiological, involving movement of existing pigments; and (ii) long-term morphological, involving synthesis or degradation of pigments (Bagnara and Hadley, 1973). Ontogeny is one of the main drivers of the observed variation in morphological melanic pigmentation patterns in insects (Booth, 1990) or fish (Gavan, 1969; Fricke, 1980). Amphibians have been far less studied in relation to melanic pigmentation variation, but spot patterns can change ontogenetically by merging, disappearing, or becoming larger (Rojas et al., 2023), as observed in *Neurergus microspilotus* newts (Vaissi et al. 2018) or *Salamandra corsica* (Beukema, 2011) yet, typically such colour changes in amphibians occur from juvenile transitions to adulthood rather than over the adult life (Hoffman & Blouin, 2000). Apart from ontogeny, explanations for changing pigmentation patterns include visual communication such as intraspecific mate recognition (Wiernasz & Kingsolver, 1992), as well as the physical properties of pigments, such as UV protection or absorption of solar energy for thermoregulation (Alho et al. 2010). Perhaps unsurprisingly, dorsal melanic pigmentation is sometimes more pronounced in montane amphibian populations (e.g., in the Alps: Nöllert & Nöllert, 1992) and has been found to increase with altitude (Behler & King, 1979) and latitude, specifically for *Rana temporaria* in northern Europe (Alho et al. 2010). Black dorsal pigmentation increased with body size for males in a high-altitude population of common frogs *R. temporaria* in northern Spain (Riobo et al., 1999) but this was estimated using a single survey and visual classification of pigmentation types, meaning actual pattern changes over time for the same individuals remain unknown. Alternatively, melanic pigmentation might be selectively neutral and maintained due to pleiotropic effects or genetic drift, with no adaptive explanation (Booth, 1990).

This study aimed to investigate pigmentation pattern change in common frogs using individual tracking over time. This set against the backdrop of the global decline in amphibian biodiversity, where a third of amphibian species are globally threatened (Stuart et al., 2004), including some previously widespread and abundant species (Petrovan & Schmidt, 2016). Amongst other causes of decline, increased UV-B radiation caused by anthropogenic ozone depletion has been identified as a significant hazard, with species breeding in exposed montane pools and temperate lakes found to be disproportionately affected by increased UV-B exposure (Beebee & Griffiths, 2005). Pigmentation patterns are also frequently employed for non-invasive monitoring to understand population size, demography and trends, including for amphibians (Ferner, 2007; Gamble et al. 2008), and pigmentation changes might invalidate capture recapture surveys that rely on such patterns to be unique in the population and stable between years. Gaining insights into pigmentation patterns as natural adaptations of amphibian species to environmental stressors (e.g. UV light), thermoregulation, predation pressure or ontogenetic changes is thus valuable from both an evolutionary understanding as well as a conservation standpoint. Our aims were therefore to: (i) study intraspecific variation in dorsal melanic pigmentation over time within a population of European common frogs that is the subject of long-term mark recapture monitoring; and (ii) determine if sex, age and spot position are significant predictors of the variation in pigmentation at the individual level.

## Materials and methods

### Focal species

*Rana temporaria* is a wide-ranging, cold tolerant terrestrial frog species distributed across Britain and much of the Western Palearctic region, from Ireland to Kazakhstan and including in some areas north of the Arctic Circle (Sillero et al. 2014). It is largely nocturnal and inhabits a variety of damp natural and semi-natural habitats including parks and gardens, and often breeds in man-made garden ponds. It is sexually dimorphic, especially during the breeding season and males are marginally smaller in length than females and, in both sexes, populations show wide variation in dorsal pigmentation (Arnold and Burton 1978). *R. temporaria* has been studied extensively and is often used to understand evolutionary adaptations in populations across its enormous distribution yet, paradoxically, dorsal pigmentation patterns remain unused as a method for mark recapture studies.

### Study Area

The studied population is located in a lowland (40m asl) suburban garden site which includes a small pond (aprox. 2m x 1m with 35cm max depth), in Stamford, Lincolnshire (52.656°N 0.484°W). The site is 100m away from large semi-natural floodplain grassland areas, and therefore unlikely to suffer significant urban heat island effects compared to dense urban areas. Eastern parts of England, where the site is located, are generally flat, drier, sunnier and warmer compared to the UK as a whole but overall the local climate is still relatively mild and wet, with a mean annual precipitation of 614 mm in 1991-2020 (England and Wales average 951 mm; UK average 1163 mm) and mean annual maximum daily temperature of 14.1 ºC (England and Wales average 13.7 ºC; UK average 12.8 ºC) (MetOffice, Wittering weather station).

### Data collection

Photographs of common frogs visiting the pond and surrounding garden were collected during August 2018 - February 2024 as part of an ongoing mark recapture monitoring project of the frog population. Photographs were taken from above to capture a full dorsal view of the frog body, using smart phone cameras (IPhone SE and 12 models). Efforts were made to maintain the same perpendicular position to the frog body axes and at the same distance from the frog to allow standardisation and reduce scaling errors. All photographs were collected without capture or handling to minimise recapture stress, potential injuries to the frogs and bias from capture avoidance. Generally, only individuals perceived to be adult size (>50 mm SVL) were included, partly to ensure these could be reliably sexed, but small numbers of 1 year old juveniles (three) were also photographed. Pigmentation patterns were used to separate and identify unique individuals using combinations of identification software (Wild-ID) and manual verifications. A single observer (SP) then visually classified all 104 recorded individuals using the detailed pigmentation chart produced by Riobo et al. (1999).

We selected a subset of 42 adult frogs, split roughly equally between males and females, by prioritising individuals with good quality, unobstructed view, full-body photographs for subsequent in depth data extraction. The sex of individual frogs was assigned using external characters such as proportional size of forearms, behaviour and nuptial colouration during the breeding season. Age was estimated based on total body size and individuals were classified as juveniles (1 year old, both sexes), young adults (2 year old males and 3 year old females) and older adults (3 or 4 or more years old depending on sex).

Digital photographs were processed using the software ImageJ version 1.53a (Schneider et al. 2012). We restricted the region of interest to the large central area between the two dorsolateral dermal plicae of the frog using the polygon selection tool and blocked the rest of the image (Figure 1). This was because areas outside the dorsolateral plicae are mainly lateral parts of the body and much more susceptible to angle distortions. We applied Gaussian Blur and the Median filter to adjust for light reflection glare wherever necessary and converted the colour image to a black and white 8-bit image and set image scale to account for differences in resolution between the pictures (Figure 1). We differentiated the dark spots from the background by using either the Auto or Manual Thresholding tool, depending on which method ensured the highest visibility for the spot selection for each picture. Finally, we used the Magic Selection tool followed by the freehand selection tool where necessary to improve accuracy of the spot selection, and then obtained area measurements of pigmentation spots from the Measure tool (Figure 1). Apart from obtaining area data, we counted manually the number of spots visible in the region of interest in each photograph.

**Figure 1.**
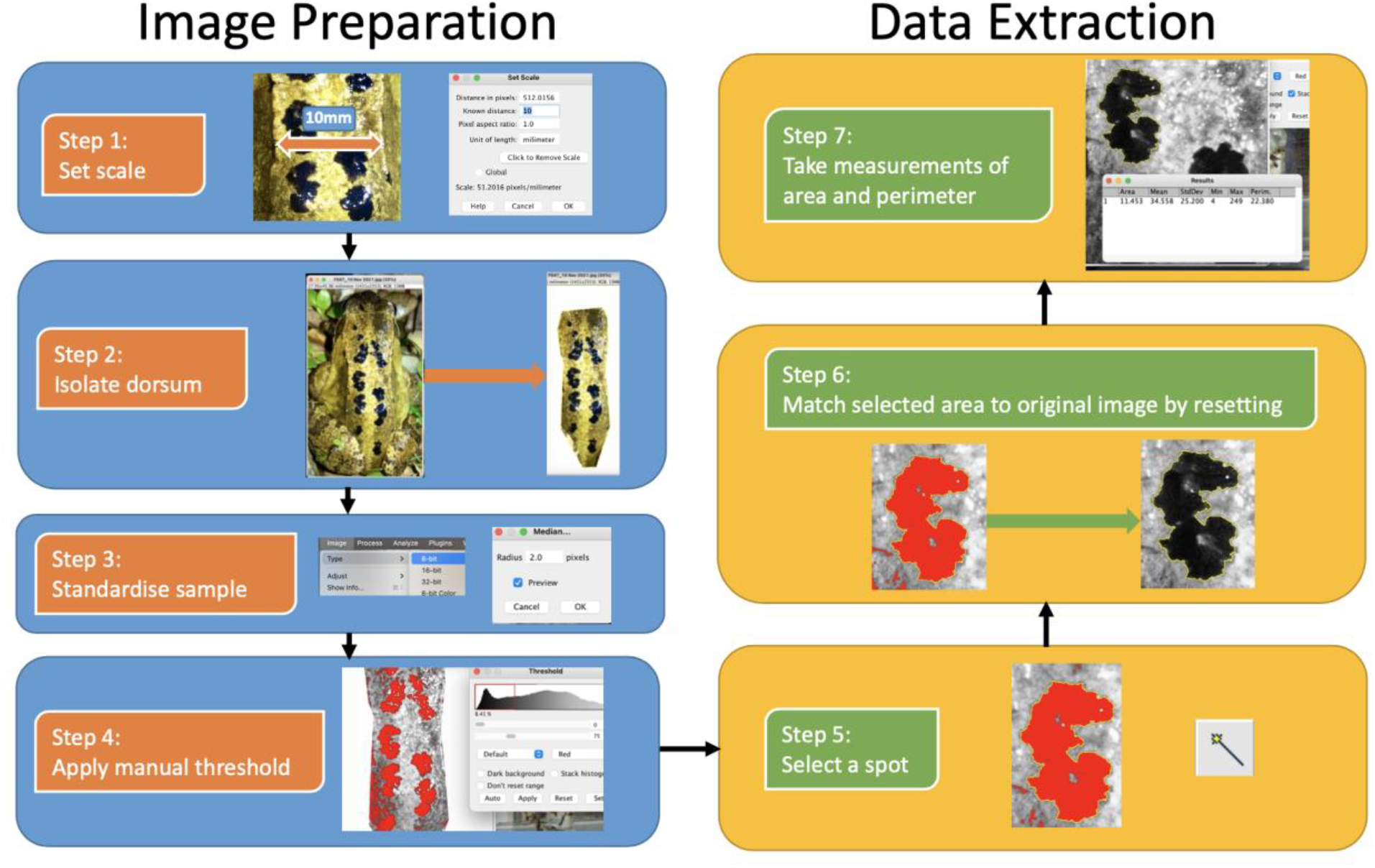
A step-by-step image preparation and data extraction framework to quantify melanin-based spot pigmentation in common frogs.

### 2.4 Data and Statistical Analyses

To quantify differences in pigmentation between sexes, we used Generalised Linear Models (GLMs) with the following formulae for two measures of pigmentation: Number of spots ∼ Sex; Total area of dorsal pigmentation ∼ Sex. For the categorical variable of sex, we set females as the intercept. For the number of spots, we used a Poisson error family because this was a count variable. For the total area of dorsal pigmentation we used a Gamma error family with a log link function because this was a positive non-zero continuous variable with a right skewed distribution (i.e., more smaller values than large ones). To quantify differences in pigmentation class between sexes, we used an ordinal regression as pigmentation class (formula: Pigmentation class ∼ Sex) is an ordinal variable ranging from 1 to 6, where 6 is most pigmented and 1 is least. For all analyses, we excluded individuals of undetermined sex.

To quantify changes in pigmentation over time for individual frogs, we used Generalised Linear Mixed Models via the glmer function of the lmer4 package in R (Bates et al., 2015) – this uses restricted maximum likelihood (REML) and the bobyqa optimizer. For our first analysis, concerning overall pigmentation changes, our response variables were the total number of spots and the total area of pigmentation on the frog dorsum. We used the timepoint (of measurement) as a fixed effect and individual identity as a random effect because some individuals were measured multiple times. The same error families were used as for the previous analyses of these response variables (i.e., Poisson for number of spots and Gamma with a log link function for pigmented area). For our second analysis to explore whether individual frogs in higher pigmentation classes were initially more pigmented to begin with, we used a simple General Linear Model with a Gaussian error family to model the response variable of the change in the pigmentation class of frogs (the difference between two pigmentation class measurements for frogs sampled at least two years apart) against the initial recorded pigmentation class and the time interval between measurements as fixed effects.

Where necessary we tested the assumptions of normality and homogeneity of variances using the DHARMa package (Hartig, 2022). We used R version 4.1.2 for all statistical analyses (R Core Team 2021).

## Results

### Differences between sexes

The total number of dorsal spots and total area of dorsal pigmentation ranged from 0 – 36 and 3.13 mm2 – 171.19 mm2 respectively (*Fig 2*). There was a wide variation range between individuals of the same sex (Table 1; Fig 2) and females on average had more spots than males but the difference was not statistically significant (p=0.077; Z=-1.77; Table 2). In terms of total area of dorsal pigmentation, males tended to be more pigmented on average (Table 1; Fig 2) but the difference was not statistically significant (p=0.364; Z=0.918; Table 2).

**Table 1.**
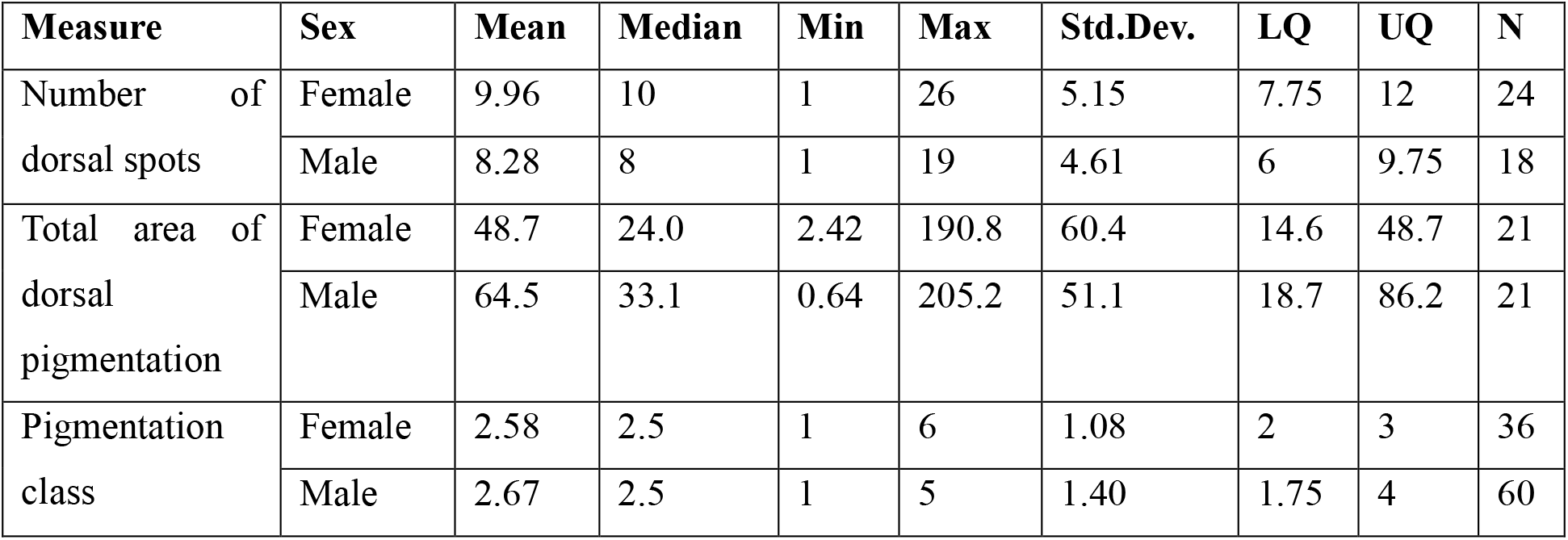
Summary statistics for the number of dorsal spots and total area of dorsal pigmentation of male and female frogs (n= 42). For individuals with repeated measurements only the most recent one was included. Frogs that were of unidentified sex were excluded. LQ = Lower Quartile; UQ = Upper Quartile.

**Table 2.** Statistics from Generalised Linear Models (GLMs) comparing number of dorsal spots and total area of dorsal pigmentation between sexes (n= 42). See methods for specifics of GLMs used. For individuals with repeated measurements only the most recent one was included. Frogs that were of unidentified sex were excluded.

| Measure | Term | Estimate | Std. Error | Z value | p-value | Pseudo-R-squared |
| --- | --- | --- | --- | --- | --- | --- |
| Number of dorsal spots | Intercept (Female) | 2.30 | 0.065 | 35.5 | 0.000 | 0.027 |
|  | Sex (Male) | -0.185 | 0.104 | -1.77 | 0.077 |  |
| Total area of dorsal pigmentation | Intercept (Female) | 3.89 | 0.217 | 17.9 | 0.000 | 0.015 |
|  | Sex (Male) | 0.282 | 0.307 | 0.918 | 0.364 |  |
|  | Sex (Male) | 0.0833 | 0.272 | 0.307 | 0.760 |  |

**Figure 2.**
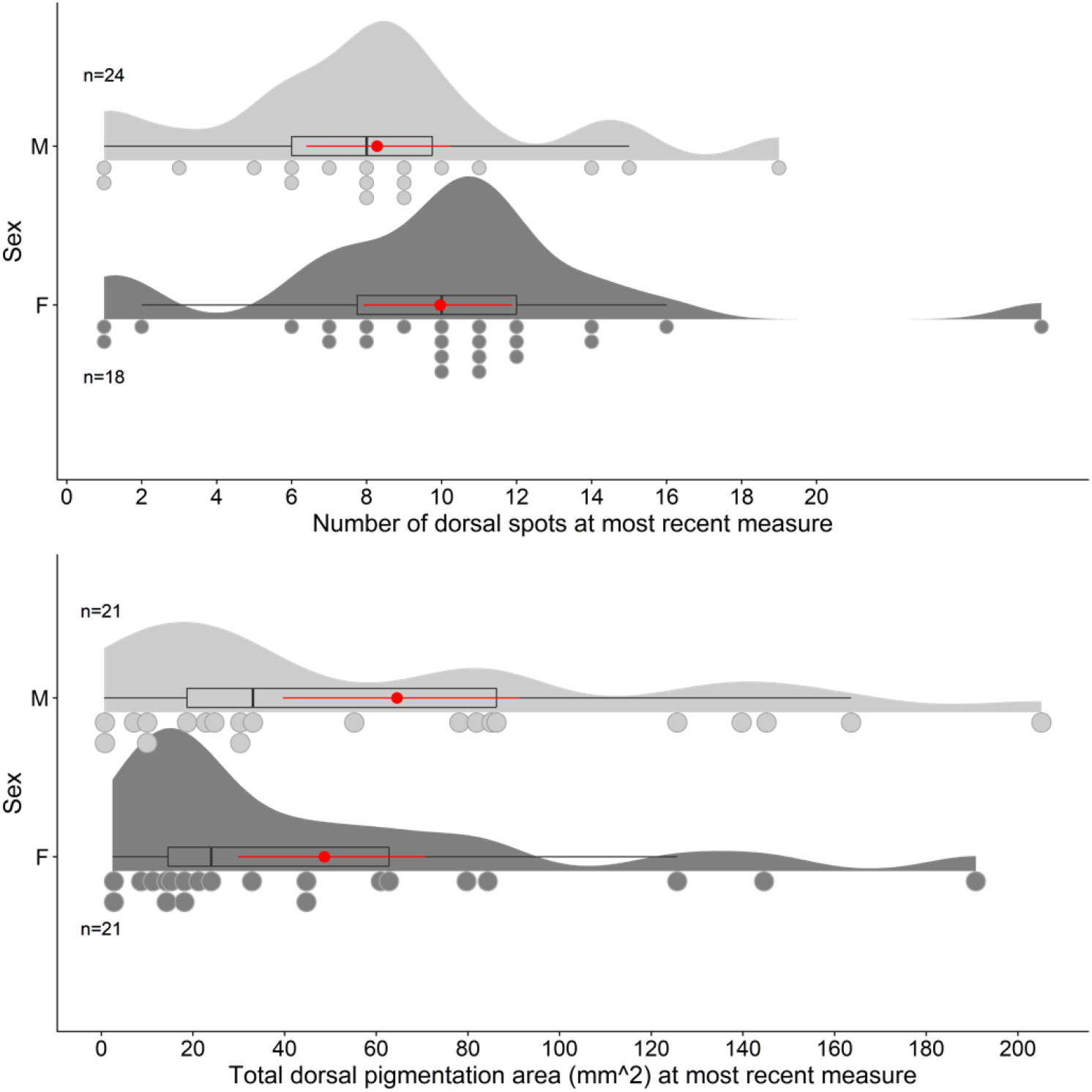
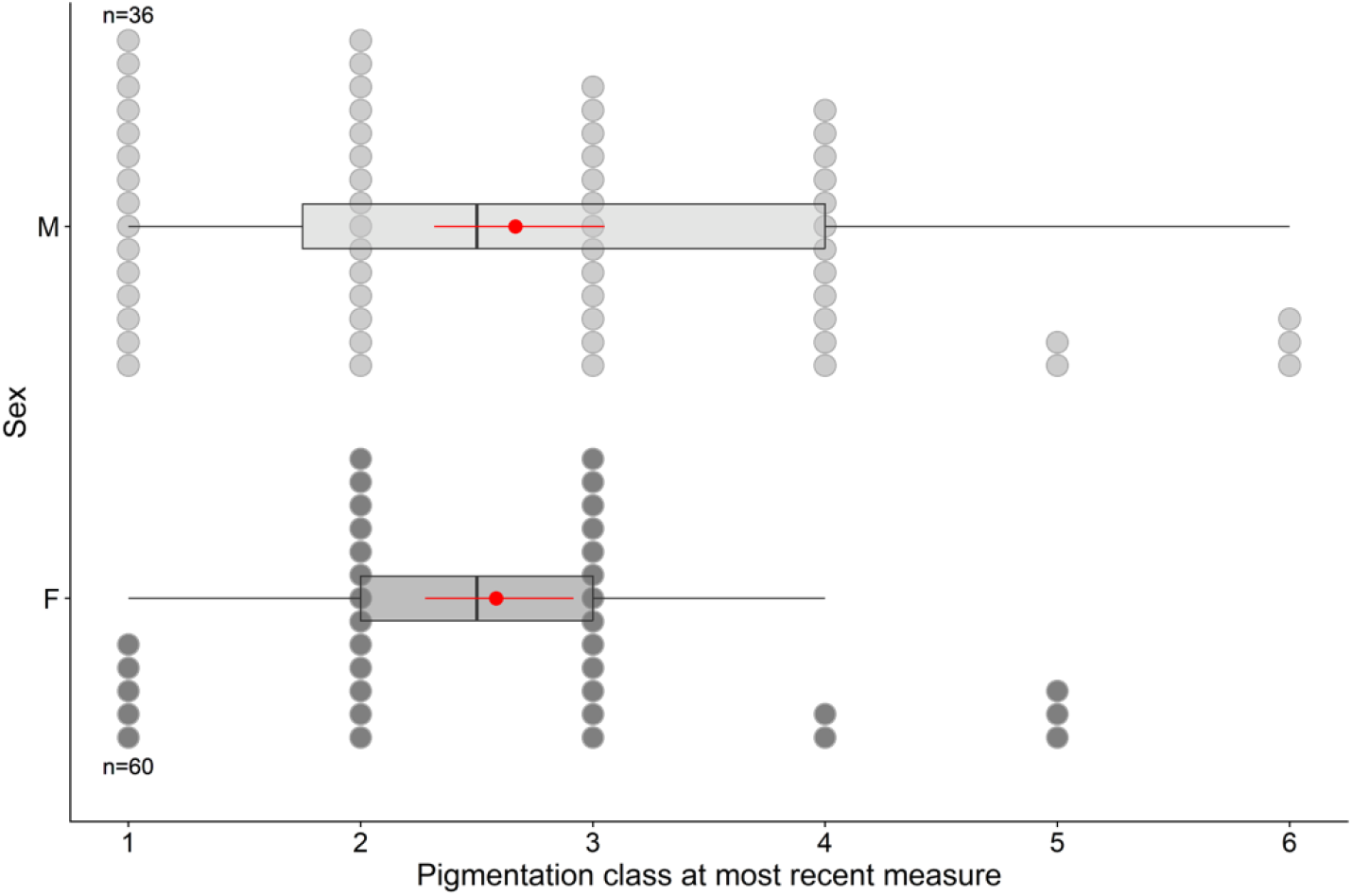
Number of dorsal spots, total area of dorsal pigmentation, and pigmentation class for males and females. For individuals with repeated measurements only the most recent one was included.

When considering pigmentation class, males were also marginally more pigmented on average (Table S1), but were not significantly more likely to be in higher pigmentation classes than females (p=0.942; t=0.0732; Odds ratio = 1.027, i.e., males were 1.02; Table 3).

**Table 3.** Statistics from an ordinal regression to compare the pigmentation classes of male and female frogs (n= 104). For individuals with repeated measurements only the most recent one was included. Frogs that were of unidentified sex were excluded. Coefficient estimates are on the log odds scale.

| Measure | Term | Estimate | Std. Error | t value | p-value |
| --- | --- | --- | --- | --- | --- |
| Pigmentation class | SexM | 0.0271 | 0.370 | 0.0732 | 0.947 |
|  | 1\|2 | -1.32 | 0.332 | -3.97 | p<0.00001 |
|  | 2\|3 | 0.0154 | 0.293 | 9.58 | 0.958 |
|  | 3\|4 | 1.23 | 0.328 | 1.77 | p<0.001 |
|  | 4\|5 | 2.41 | 0.434 | 2.64 | p<0.00001 |
|  | 5\|6 | 3.45 | 0.629 | 4.20 | p<0.00001 |

### Changes in the number of spots, area, and pigmentation class over time

There was a negligible change in the number of spots over time (p=0.091; t=-1.69; Table 4) and most of the variation explained by the model occurred between individual frogs (see large conditional R-squared value (variation explained by Frog ID) versus small marginal R-squared value (variation explained by Time-point; Table 4). On average, the number of spots declined by 0.000176 spots per day, or 0.064 spots per year, with spot merging as the most likely underlying mechanism – as observed in Figures 4-6.

**Table 4.**
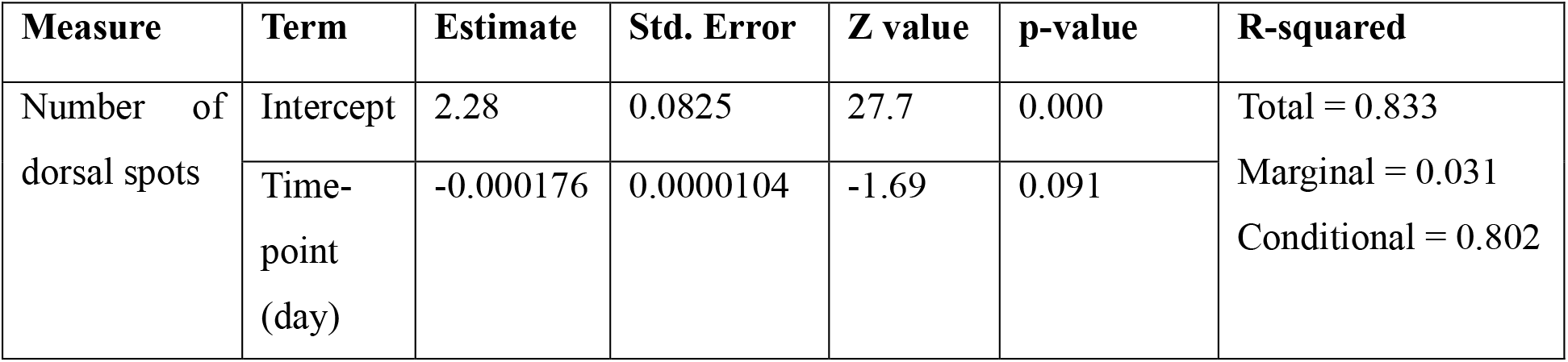
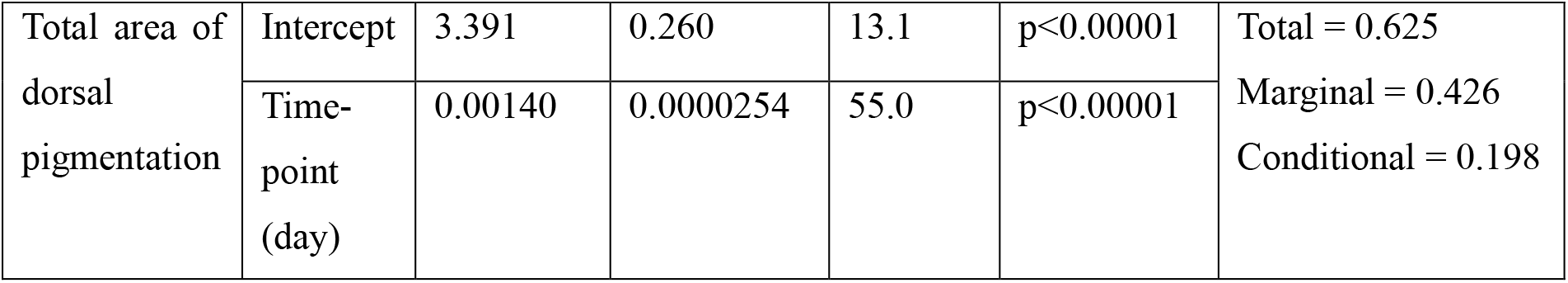
Statistics from Generalised Linear Mixed Models (GLMMs) analysing the change in the number of dorsal spots and total area of dorsal pigmentation over time. See methods for specifics of GLMMs used – a random effect was applied for frog ID due to repeated measures. Frogs that were of unidentified sex were excluded. Total, marginal and conditional R-squared values are presented.

Total dorsal pigmentation area varied widely between individuals but increased significantly over time (p<0.00001; Z = 55.0; Table 4; Fig. 3) by 1.0014 times per day (0.14% per day) on average (∼1.67 times increase in a year, 6.67 times in 4 years or 1460 days). There was no evidence of total pigmentation area reaching an asymptote.

**Figure 3.**
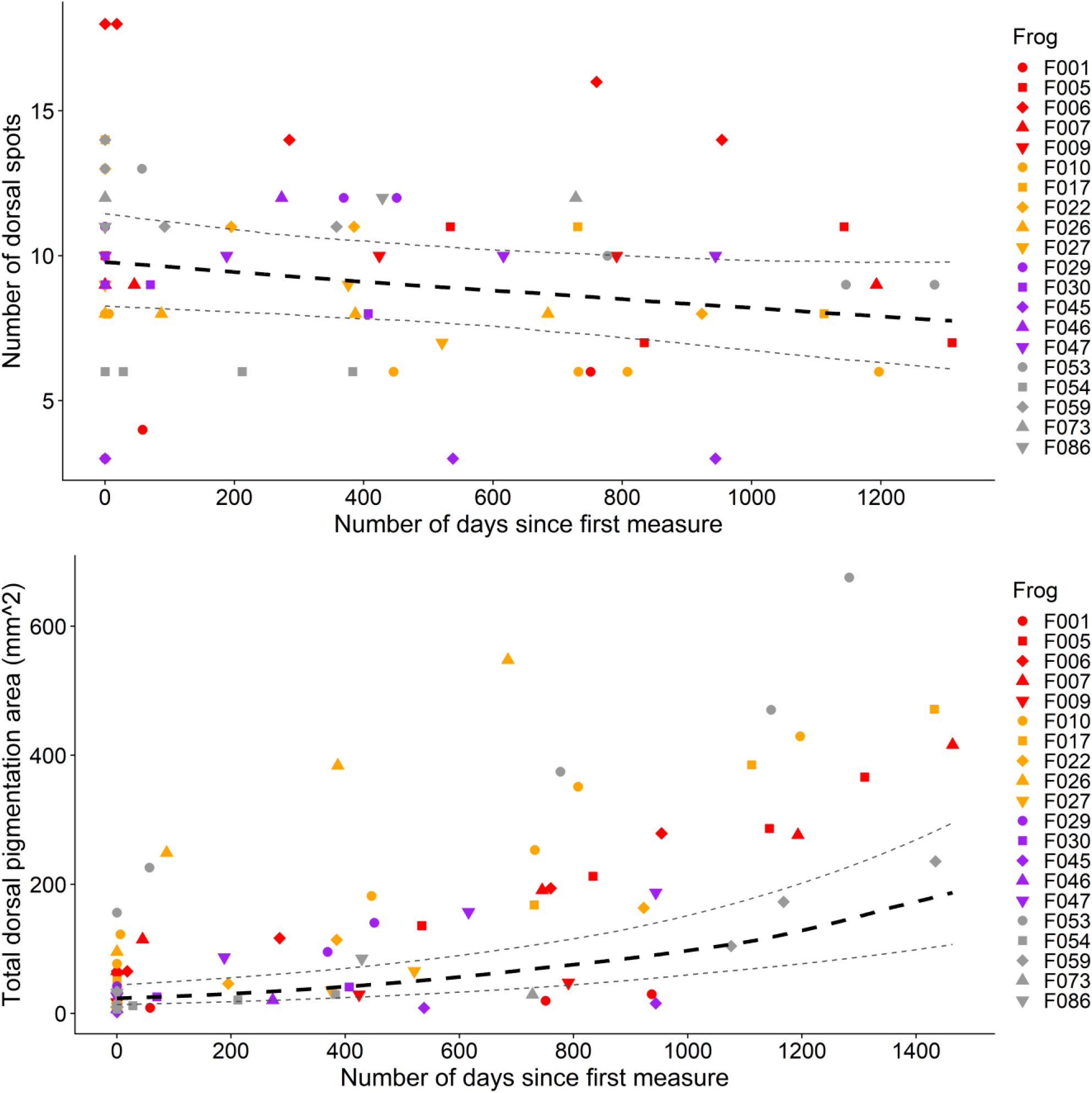
The number of spots and total dorsal pigmentation area of individual frogs over time. Black dotted lines are based on Generalised Linear Mixed Models (GLMMs) above (Table 3) with bootstrapped prediction intervals. Datapoints for individual frogs are shown with different coloured symbols.

**Figure 4.**
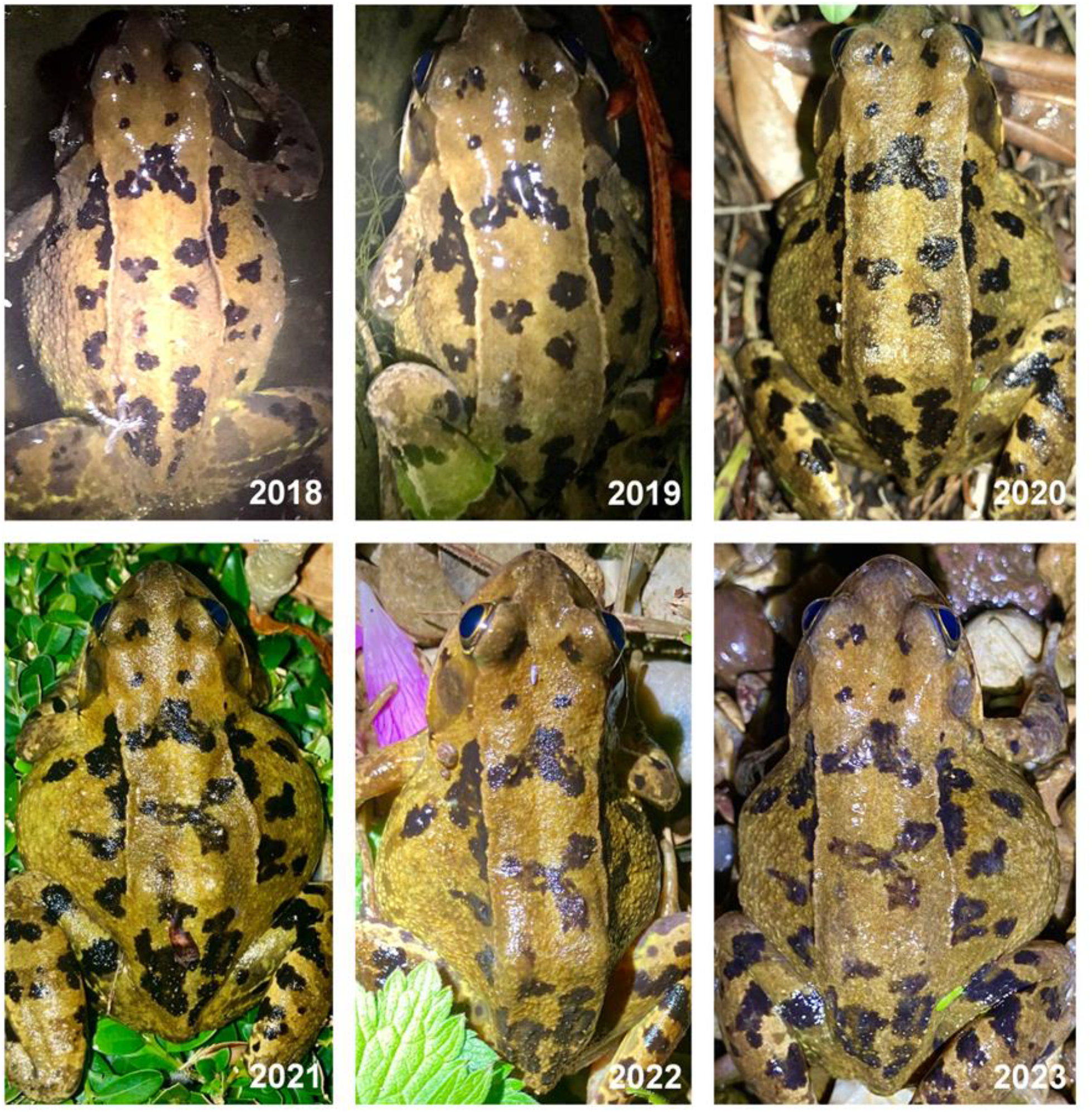
Multi-year recaptures and pigmentation pattern changes in female F005. Fusion of the three central dorsal spots is noticeable after 2020.

**Figure 5.**
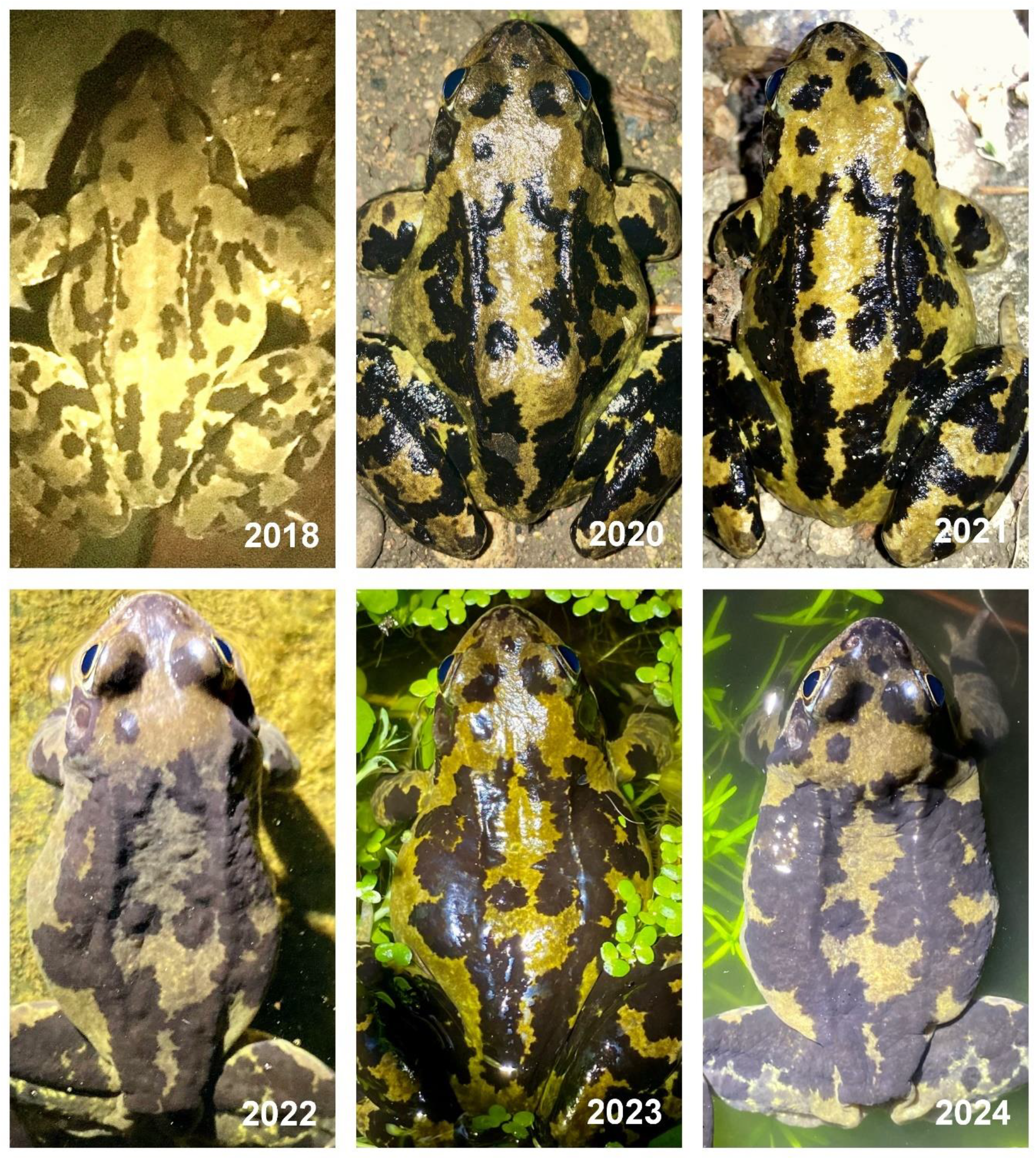
Multi-year recaptures and pigmentation changes in male F053, one of the individuals with the highest increase in pigmentation area between years.

**Figure 6.**
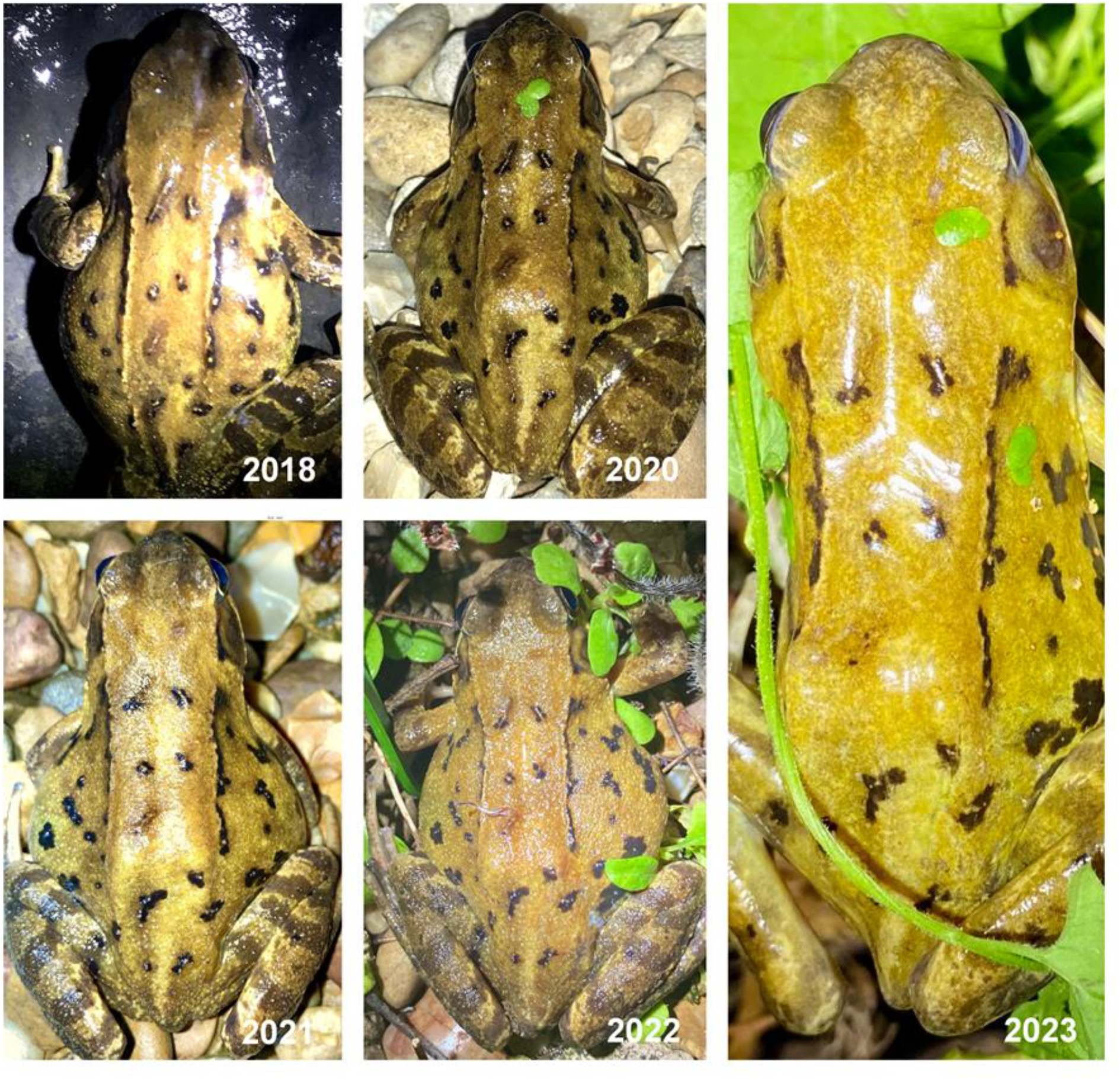
Multi-year recaptures and pigmentation changes in female F009, one of the individuals with the lowest change in pigmentation area between years.

However, individual variability was clearly apparent (Fig.3; see conditional R-squared value in Table 4), with several individuals seeing only neglible change in total dorsal pigmentation area. This seemed, at least in part, related to the starting area of spots, as frogs that were first recorded in a higher pigmentation class (specifically class 4; p=0.023; t=2.35; Table 5) increased significantly more in pigmentation class than frogs in class 1, whilst accounting for the length of the interval between measurements (greater increases in pigmentation class were observed over longer time intervals between measurements (p<0.0001, t=4.72; Table 5)).

**Table 5.** Statistics from General Linear Model analysing the change in pigmentation class in relation to the first measured pigmentation class for that individual frog and the time interval between measurements. See methods for specifics of model used. Frogs that were of unidentified sex were excluded.

| Measure | Term | Estimate | Std. Error | z/t value | p-value | Adjusted R-squared |
| --- | --- | --- | --- | --- | --- | --- |
| Change in pigmentation class | Intercept | -0.0835 | 0.222 | -0.375 | 0.710 | 0.361 |
|  | Starting Class: 2 | 0.101 | 0.250 | 0.402 | 0.690 |  |
|  | Starting Class: 3 | 0.0822 | 0.235 | 0.349 | 0.729 |  |
|  | Starting Class: 4 | 1.17 | 0.496 | 2.35 | 0.023 |  |
| | Time interval (days) | 0.001 | 0.000 | 4.72 | $p<0.0001$ | |

## Discussion

Using a novel and easy to replicate method, this study shows how pigmentation patterns change (or not) over time for the same frog individuals tracked via multiannual surveys, but also that individuals from the same population of *R. temporaria* had substantial variability in the degrees of dorsal melanin-based pigmentation. Pigmentation patterns ranged from no spots in the region of interest (observable in two adult individuals, one male, one female) to dozens of spots and a large melanistic pigmentation for several other individuals. Similar to previous findings, males tended to be darker on average while females tended to have more spots but, overall, in our dataset neither were significantly different in statistical terms. While there seems to be some general support for the darker male pigmentation, the variation between sexes is likely influenced by the age structure of the population as males might have longer life expectancy and thus are more likely to reach high degree of pigmentation, as well as baseline pigmentation levels in young adults.

Overall, we found that the total melanic area increased over time, and this was particularly apparent in some individuals, both males and females. Applying the visual classification system by Riobo et al. (1999), four individuals in our study increased their pigmentation area sufficiently to change from class 3 to the highest classes, 5-6, in the space of four years. However, while the increasing pigmentation trend was significant overall (in terms of area), several males and females showed virtually no pigmentation change across the years, and this appeared particularly the case for individual with low initial degree of melanin-based pigmentation. In essence, many strongly pigmented frogs become progressively darker with age while frogs with few and small spots as young adults were unlikely to darken significantly and most likely remained unchanged, even as old adults. This is in contradiction to previous findings which found a close correlation between body size as a proxy for age and pigmentation degree (Riobo et al., 1999). Despite the smaller sample size of individuals considered, we demonstrate a clear of progressive melanism for several adult frogs of both sexes. The implication is that the pigmentation degree is not a valid proxy for age approximation, and especially for individuals with relatively low levels of pigmentation. Understanding the genetic, evolutionary and morphological basis for such pigmentation differences could be an important focus for future research, especially as melanophore proliferation has wider scientific relevance, including immunity, hormonal and oncological changes, and there are even important parallels between frog skin and human skin (Haslam et al. 2014).

Although the darkest individuals were mainly males, overall males and females had a similar proportion of dorsal pigmentation yet this could be influenced by different survival rates between sexes and the high pigmentation variability within sex in this population. Dorsal melanism is plausibly a trait under some degree of sexual selection interacting with ontogenetic factors but could equally well be result of some aspect of natural selection acting on males more than females, for instance related to the earlier presence at the pond by males in January-February when temperatures are often still under 5C. However, this remains untested and requires additional confirmation.

This ontogenetic variation, with most frogs becoming progressively darker could be associated with reproductive status (e.g., ontogenetic dichromatism: Lambert et al., 2017), where the sexes diverge in melanism patterns during development. Such divergence facilitates conspecific identification in terms of (i) sex (male vs female) and (ii) reproductive status (adult vs juvenile). Distinct melanism between sexes could enable males to differentiate between other males and female mates and this might be particularly significant for the time constraints in explosive breeding events in *R. temporaria*, where rapid sex-identification of conspecifics is essential for higher breeding success. However, if this were the case we would expect a strong divergence in melanism between males and females favoured by directional selection, yet this was not the case in our results. Secondly, sexually mature individuals have been observed to have a distinct colouration compared to juveniles in order to prevent occurrences of mating attempts with prereproductives (Booth, 1990) and only reproductive-age males of this species call during the breeding period.

Many studies have also demonstrated that the degree of melanism is associated with body condition - traits such as body mass or immunity that reflect better health, viability or fertility - across taxa (birds: Hadfield & Owens, 2006; fish: Wedekind et al., 2008; Marie-Orleach et al., 2014). This links to the ‘Handicap principle’, proposed by Zahavi (1975), which posits that secondary sexual traits that are costly to produce serve as honest signals that can be used for mate choice. Contrary to classic sexual selection theory, with males having costly ornaments to impress the females, our results indicate that males and females had a similar degree of pigmentation. This makes sense in the light of the strong association found between female body size and fecundity in explosively breeding species, such as *R. temporaria* (Nali et al., 2014), and that male mate choice in anurans can arise under conditions where size-related fecundity variation exists (Krupa, 1995). Therefore, the higher dorsal pigmentation in some males could signal higher fecundity and nutritional status to attract mating females or to deter competition with other males. However, a cursory analysis of the observations of males in amplexus at this site did not indicate any higher rates of success for more pigmented males and in fact some appeared to be significantly less successful compared to other, less pigmented males. Further evidence is required to clarify the relationship between amphibian colouration and nutritional status but darker individuals in this study were not necessarily the largest. Also, the literature concerning the influence of male mate choice in anurans remains inconclusive, so it is difficult to draw conclusions regarding the influence of sexual selection on the degree of melanism in frogs. The mechanisms that promote color polymorphisms in nocturnal species, such as common frogs, remain poorly explored compared to diurnal species but there is a possible role of sexual selection and/or differential predation pressures on maintaining color morphs in nocturnal species (Aguilar et al. 2023).

Identification of individual animals is a key part of many field studies. Marked individuals can provide a wealth of information on population dynamics, geographic range, animal behaviour and many other demographic, reproductive or ethological characteristics (Silvy, 2020). While a broad range of invasive identification methods are used, such as toe clipping, PIT-tagging or implanting elastomers, they can induce significant stress, affecting the survival and behaviour of the animal (Antwis et al., 2014) and (ii) can be logistically difficult and expensive, especially for longitudinal studies (Donnelly et al. 1994). The results of our study demonstrate significant individual differences in dorsal pigmentation, which, even without any established adaptive significance, has the potential to be used as a characteristic for non-invasive identification of frog species. The fact that the number of spots was largely unchanged over time is important as an indicator of the general stability of such melanin-based spot pigmentation in this species and especially as none of the tracked spots disappeared. This marks an important difference from other amphibian species where spots were found to disappear or migrate to new regions (e.g. Vaissi et al., 2018) but also highlights the apparent permanence of melanin-based pigmentation in this species. Pattern stability combined with the pattern complexity are also indicators of the value of using spot pigmentation pattern as an individual identification and mark recapture tool for this species. The combination of shape, size and position of a multitude of melanin-based pigmentation spots allows for almost infinite numbers of unique combinations between individuals of this species, making pattern recognition an obvious monitoring tool in future studies. However, this is not absolute as two individuals in this study had no spots at all in the dorsal region of interest although they had recognisable melanin-based spots on the limbs. Populations where this absence of melanin-based patterns in the dorsal area as a form of hypopigmentation is relatively common would make this identification method error prone and unfeasible. We recommend initial pilot surveys and even the use of the simple pigmentation classification to verify the pattern variability and the presence of individuals without pigmentation spots in the population as a useful precondition to undertaking mark recapture surveys. Future studies could link pigmentation and estimated age with other methods such as skeletochronology to get additional confirmation. However, it is unclear how robust would estimations of skeletochronology be given that frogs in this population were shown to be active throughout the entire year (Petrovan et al. in review).

Methods of pigmentation analysis for wildlife identification purposes are increasing in popularity, especially for amphibians (Carafa & Biondi, 2004; Gamble et al., 2008; Mettouris et al., 2016), but are currently primed for a dramatic leap forward in terms of volume, speed and efficiency with the incorporation of Machine Learning and AI approaches. Increasing our understanding of the factors affecting such patterns and pigmentation on animals would enable us to make assessments on the stability of such pigmentation and identify individuals with a higher degree of accuracy. The accessible methodology of our study was designed to verify the potential applicability as a citizen science tool for large-scale data collection on this widespread species. First, collection methods such as mobile phone photographs are easy-to-use for members of the general public and allow centralised separate-stage analysis to facilitate replicability. Secondly, citizen scientists greatly increase the geographic reach of field work and can collect data on private land that would otherwise be difficult and expensive to access at scale. Thirdly, public engagement is at the core of any biodiversity crisis – getting members of the public to participate in science is one of the best ways to get them to understand and appreciate science and biodiversity. Citizen science, combined with automation and machine learning, could dramatically increase the scope of such data collection for amphibians and provide in-depth monitoring and evolutionary adaptation data that could assist us in better protecting them.

## Acknowledgements

Andrew Balmford contributed significantly during the project supervision. We thank Roger Downie and Emilia Santos for useful comments and discussions on the manuscript draft.

## Author Contributions

*SP* conceived the ideas, collected the data and designed the methodology; *SS* and *CL* analysed the image database and extracted the measurements; *AC* led the statistical analysis; SP led the writing of the manuscript. All authors contributed to the drafts and gave final approval for publication.

## Competing Interests

None

**Table S1.**
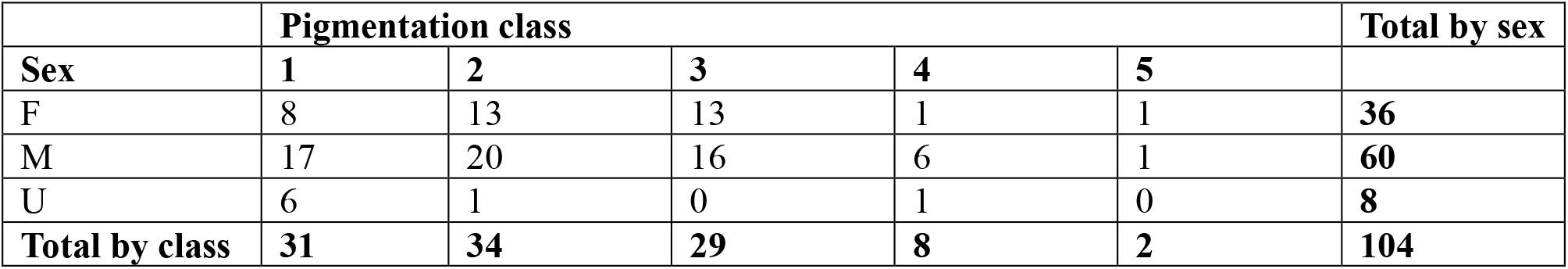
Summary counts of frogs by the visual pigmentation classification.

## References

Aguilar, P., Pérez i de Lanuza, G., Martínez‐Gil, H., Dajčman, U., Simčič, T., Pinho, C., Žagar, A. and Megía‐Palma, R. (2023). Color morphs of the fire salamander are discriminated at night by conspecifics and predators. Journal of Zoology. 10.1111/jzo.13131

Alho, J. S., Herczeg, G., Söderman, F., Laurila, A., Jönsson, K. I., & Merilä, J. (2010). Increasing melanism along a latitudinal gradient in a widespread amphibian: local adaptation, ontogenic or environmental plasticity?. BMC Evolutionary Biology, 10(1), 1–9.

Antwis, R. E., Garcia, G., Fidgett, A. L., & Preziosi, R. F. (2014). Tagging frogs with passive integrated transponders causes disruption of the cutaneous bacterial community and proliferation of opportunistic fungi. Applied and Environmental Microbiology, 80(15), 4779–4784. 10.1128/AEM.01175-14

Arnold, N., & Ovenden, D. (1978). A field guide to the reptiles and amphibians of Britain and Europe. Collins, London.

Bagnara, J. T., & Hadley, M. E. (1973). Chromatophores and color change: The comparative physiology of animal pigmentation. Prentice-Hall.

Beebee, T. J. C., & Griffiths, R. A. (2005). The amphibian decline crisis: A watershed for conservation biology? Biological Conservation, 125(3), 271–285. 10.1016/j.biocon.2005.04.009

Behler, J. L., & King, F. W. (1979). The Audubon Society field guide to North American reptiles and amphibians (A Chanticleer Press ed). Knopf : Distributed by Random House.

Beukema, W. (2011). Ontogenetic pattern change in amphibians: the case of Salamandra corsica. Acta Herpetologica, 6(2), 169–174.

Booth, C. L. (1990). Evolutionary significance of ontogenetic colour change in animals. Biological Journal of the Linnean Society, 40(2), 125–163. 10.1111/j.1095-8312.1990.tb01973.x

Carafa, M., & Biondi, M. (2004). Application of a method for individual photographic identification during a study on Salamandra salamandra gigliolii in central Italy. Italian Journal of Zoology, 71(Sup2), 181–184. 10.1080/11250000409356631

Donnelly, M. A., Guyer, C., Juterbock, E. J., & Alford, R. A. (1994). Techniques for marking amphibians. Measuring and monitoring biological diversity: standard methods for amphibians.

Ducrest, A.-L., Ursenbacher, S., Golay, P., Monney, J.-C., Mebert, K., Roulin, A., & Dubey, S. (2014). Pro-opiomelanocortin gene and melanin-based colour polymorphism in a reptile: Colour Polymorphism. Biological Journal of the Linnean Society, 111(1), 160–168. 10.1111/bij.12182

Ethier, J., Gasse, M., Lake, K., Jones, B. C., Evenden, M. L., & Despland, E. (2015). The costs of colour: Plasticity of melanin pigmentation in an outbreaking polymorphic forest moth. Entomologia Experimentalis et Applicata, 154(3), 242–250. 10.1111/eea.12275

Ferner, J. W. (2007). A review of marking and individual recognition techniques for amphibians and reptiles. Herpetological Circular 9. Society for the Study of Amphibians and Reptiles. Salt Lake City, UT.

Fricke, H. W. (1980). Juvenile-adult Colour Patterns and Coexistence in the Territorial Coral Reef Fish Pomacanthus imperator. Marine Ecology, 1(2), 133–141. 10.1111/j.1439-0485.1980.tb00215.x

Fulgione, D., Trapanese, M., Maselli, V., Rippa, D., Itri, F., Avallone, B., Van Damme, R., Monti, D.M. and Raia, P., 2014. Seeing through the skin: dermal light sensitivity provides cryptism in moorish gecko. Journal of Zoology, 294(2), pp.122–128. 10.1111/jzo.12159

Gamble, L., Ravela, S., & McGarigal, K. (2008). Multi-scale features for identifying individuals in large biological databases: An application of pattern recognition technology to the marbled salamander Ambystoma opacum: Identifying individual marbled salamanders. Journal of Applied Ecology, 45(1), 170–180. 10.1111/j.1365-2664.2007.01368.x

Gangoso, L., Grande, J. M., Ducrest, A.-L., Figuerola, J., Bortolotti, G. R., Andrés, J. A., & Roulin, A. (2011). MC1R-dependent, melanin-based colour polymorphism is associated with cell-mediated response in the Eleonora’s falcon: MC1R-based colouration and inflammatory response. Journal of Evolutionary Biology, 24(9), 2055–2063. 10.1111/j.1420-9101.2011.02336.x

Gavan, J. A. (1969).: A Handbook of Living Primates: Morphology, Ecology and Behaviour of Nonhuman Primates. J. R. Napier, P. H. Napier. American Anthropologist, 71(2), 357–358. 10.1525/aa.1969.71.2.02a00550

Hadfield, J. D., & Owens, I. P. F. (2006). Strong environmental determination of a carotenoid-based plumage trait is not mediated by carotenoid availability. Journal of Evolutionary Biology, 19(4), 1104–1114. 10.1111/j.1420-9101.2006.01095.x

Hartig, F. (2022). DHARMa: Residual Diagnostics for Hierarchical (Multi-Level / Mixed) Regression Models. R package version 0.4.6. https://CRAN.R-project.org/package=DHARMa

Haslam, I.S., Roubos, E.W., Mangoni, M.L., Yoshizato, K., Vaudry, H., Kloepper, J.E., Pattwell, D.M., Maderson, P.F. and Paus, R. (2014). From frog integument to human skin: dermatological perspectives from frog skin biology. Biological Reviews, 89(3), 618–655.

Hoffman, E. A., & Blouin, M. S. (2000). A review of colour and pattern polymorphisms in anurans. Biological Journal of the Linnean Society, 70(4), 633–665.

Jacquin, L., Lenouvel, P., Haussy, C., Ducatez, S., & Gasparini, J. (2011). Melanin-based coloration is related to parasite intensity and cellular immune response in an urban free living bird: The feral pigeon Columba livia. Journal of Avian Biology, 42(1), 11–15. 10.1111/j.1600-048X.2010.05120.x

Karell, P., Kohonen, K., & Koskenpato, K. (2021). Specialist predation covaries with colour polymorphism in tawny owls. Behavioral Ecology and Sociobiology, 75(3), 45. 10.1007/s00265-021-02986-6

King, R. B. (1987). Color pattern polymorphism in the Lake Erie water snake, Nerodia sipedon insularum. Evolution, 41(2), 241–255. 10.1111/j.1558-5646.1987.tb05794.x

Kittilsen, S., Schjolden, J., Beitnes-Johansen, I., Shaw, J. C., Pottinger, T. G., Sørensen, C., Braastad, B. O., Bakken, M., & Øverli, Ø. (2009). Melanin-based skin spots reflect stress responsiveness in salmonid fish. Hormones and Behavior, 56(3), 292–298. 10.1016/j.yhbeh.2009.06.006

Krupa, J. J. (1995). How Likely Is Male Mate Choice among Anurans? Behaviour, 132(9/10), 643–664. JSTOR.

Lambert, M., Carlson, B., Smylie, M., & Swierk, L. (2017). Ontogeny of Sexual Dichromatism in the Explosively Breeding Wood Frog. Herpetological Conservation and Biology, 12, 447–456.

Marie-Orleach, L., Roussel, J.-M., Bugeon, J., Tremblay, J., Ombredane, D., & Evanno, G. (2014). Melanin-based coloration of sneaker male Atlantic salmon is linked to viability and emergence timing of their offspring: Good Genes in Atlantic Salmon Sneakers? Biological Journal of the Linnean Society, 111(1), 126–135. 10.1111/bij.12187

Mettouris, O., Megremis, G., & Giokas, S. (2016). A newt does not change its spots: Using pattern mapping for the identification of individuals in large populations of newt species. Ecological Research, 31(3), 483–489. 10.1007/s11284-016-1346-y

Nali, R. C., Zamudio, K. R., Haddad, C. F. B., & Prado, C. P. A. (2014). Size-Dependent Selective Mechanisms on Males and Females and the Evolution of Sexual Size Dimorphism in Frogs. The American Naturalist, 184(6), 727–740. 10.1086/678455

Nöllert, A., & Nöllert, C. (1992). Die Amphibien Europas: Bestimmung, Gefährdung, Schutz. Franckh-Kosmos Patrelle, C., Hjernquist, M. B., Laurila, A., Söderman, F., & Merilä, J. (2012). Sex differences in age structure, growth rate and body size of common frogs Rana temporaria in the subarctic. Polar Biology, 35(10), 1505–1513. 10.1007/s00300-012-1190-7

Petrovan, S. O., & Schmidt, B. R. (2016). Volunteer Conservation Action Data Reveals Large-Scale and Long-Term Negative Population Trends of a Widespread Amphibian, the Common Toad (Bufo bufo). PLOS ONE, 11(10), e0161943. 10.1371/journal.pone.0161943

Riobo, A., Rey, J., Puente, M., Miramontes, C., & Vences, M. (1999). Ontogenetic increase of black dorsal pattern in Rana temporaria. BULLETIN-BRITISH HERPETOLOGICAL SOCIETY, 70, 1–6.

Rojas, B., Lawrence, J. P., & Márquez, R. (2023). Amphibian Coloration: Proximate Mechanisms, Function, and Evolution. In Evolutionary Ecology of Amphibians (pp. 219–258). CRC Press. 10.1201/9781003093312

Roulin, A. (2014). Melanin-based colour polymorphism responding to climate change. Global Change Biology, 20(11), 3344–3350. 10.1111/gcb.12594

Roulin, A. (2016). Condition-dependence, pleiotropy and the handicap principle of sexual selection in melanin-based colouration: Melanin and sexual selection. Biological Reviews, 91(2), 328–348. 10.1111/brv.12171

Schneider, C. A., Rasband, W. S., & Eliceiri, K. W. (2012). NIH Image to ImageJ: 25 years of image analysis. Nature Methods, 9(7), 671–675. doi:10.1038/nmeth.2089

Sillero, N., Campos, J., Bonardi, A., Corti, C., Creemers, R., Crochet, P., Crnobrnja Isailović, J., Denoël, M., Ficetola, G. F., Gonçalves, J., Kuzmin, S., Lymberakis, P., de Pous, P., Rodríguez, A., Sindaco, R., Speybroeck, J., Toxopeus, B., Vieites, D. R., & Vences, M. (2014). Updated distribution and biogeography of amphibians and reptiles of Europe. Amphibia-Reptilia, 35(1), 1–31. 10.1163/15685381-00002935

Silvy, N. J. (2020). The Wildlife Techniques Manual: Volume 1: Research. Volume 2: Management. (Issue v. 1). Johns Hopkins University Press. https://books.google.co.uk/books?id=35zwDwAAQBAJ

Stuart, S. N., Chanson, J. S., Cox, N. A., Young, B. E., Rodrigues, A. S. L., Fischman, D. L., & Waller, R. W. (2004). Status and Trends of Amphibian Declines and Extinctions Worldwide. Science, 306(5702), 1783–1786. 10.1126/science.1103538

Turner, J. R. (1977). Butterfly mimicry: the genetical evolution of an adaptation. Evolutionary biology, 10, 63–206.

Zahavi, A. (1975). Mate selection—a selection for a handicap. Journal of theoretical Biology, 53(1), 205–214.

Vaissi, S., Parto, P., & Sharifi, M. (2018). Ontogenetic changes in spot configuration (numbers, circularity, size and asymmetry) and lateral line in Neurergus microspilotus (Caudata: Salamandridae). Acta Zoologica, 99(1), 9–19. 10.1111/azo.12187

Vences, M., Galán, P., Vieites, D. R., Puente, M., Oetter, K., & Wanke, S. (2002). Field body temperatures and heating rates in a montane frog population: The importance of black dorsal pattern for thermoregulation. Annales Zoologici Fennici, 39(3), 209–220. JSTOR.

Wiernasz, D. C., & Kingsolver, J. G. (1992). Wing melanin pattern mediates species recognition in Pieris occidentalis. Animal Behaviour, 43(1), 89–94. 10.1016/S0003-3472(05)80074-0

